# The 3’ UTR of *vigR* is required for virulence in *Staphylococcus aureus* and has expanded through STAR sequence repeat insertions

**DOI:** 10.1101/2023.05.16.541063

**Authors:** Daniel G. Mediati, William Dan, David Lalaouna, Hue Dinh, Alaska Pokhrel, Timothy P. Stinear, Amy K. Cain, Jai J. Tree

**Author notes:** Corresponding author: Jai J. Tree.

## Abstract

*Staphylococcus aureus* is an adaptable human pathogen causing life-threatening endocarditis and bacteraemia. Methicillin-resistant *S. aureus* (MRSA) is alarmingly common, and treatment is confined to last-line antibiotics. Vancomycin is the treatment of choice for MRSA bacteraemia and vancomycin treatment failure is often associated with vancomycin-intermediate *S. aureus* strains termed VISA. The regulatory 3’ UTR of *vigR* mRNA contributes to vancomycin tolerance in the clinical VISA isolate JKD6008 and upregulates the lytic transglycosylase IsaA. Using MS2-affinity purification coupled with RNA sequencing (MAPS), we find that the *vigR* 3’ UTR also interacts with mRNAs involved in carbon metabolism, amino acid biogenesis, cell wall biogenesis, and virulence. The *vigR* 3’ UTR was found to repress *dapE*, a succinyl-diaminopimelate desuccinylase required for lysine and cell wall peptidoglycan synthesis, suggesting a broader role in controlling cell wall metabolism and vancomycin tolerance. Deletion of the *vigR* 3’ UTR increased VISA virulence in a wax moth larvae model, and we find that an *isaA* mutant is completely attenuated in the larvae model. Sequence and structural analysis of the *vigR* 3’ UTR indicates that the UTR has expanded through the acquisition of *Staphylococcus aureus* repeat insertions (STAR repeats) that partly contribute sequence for the *isaA* interaction seed and may functionalise the 3’ UTR. Our findings reveal an extended regulatory network for *vigR*, uncovering a novel mechanism of regulation of cell wall metabolism and virulence in a clinical *S. aureus* isolate.

## INTRODUCTION

*Staphylococcus aureus* is an adaptable human pathogen and a major cause of life-threatening endocarditis and bacteraemia (Tong et al., 2015). *S. aureus* colonises almost every site in the human body and is increasingly associated with colonisation of medical implants. Treatment has been complicated by the emergence of antibiotic-resistant strains, particularly methicillin-resistant *S. aureus* (MRSA) where treatment is limited to last-line antibiotics. The cell wall-targeting glycopeptide antibiotic vancomycin is the treatment of choice for MRSA bacteraemia. However, vancomycin treatment failure is increasingly common and attributed to MRSA isolates (up to 13%) with intermediate vancomycin resistance (4-8μg/mL) termed vancomycin-intermediate *S. aureus* (VISA) (Howden et al., 2010). VISA isolates have thicker bacterial cell walls that likely limits the permeability of vancomycin to the division septum where binding to cell wall precursors occurs. Single nucleotide polymorphisms in transcriptional regulators have been reported (Howden et al., 2010), suggesting that loss-of-function mutations and changes in gene regulation promote vancomycin tolerance.

Regulatory non-coding RNA (ncRNA) are gene regulators that typically range from 50-500 nt and control the expression of target messenger RNA (mRNA) through direct base-pairing. Interactions between ncRNA-mRNA can promote or inhibit degradation by cellular ribonucleases such as RNases that are often associated with the RNA degradosome (Bandyra et al., 2012; Papenfort et al., 2013). The canonical pathway of gene regulation involves occluding the ribosomal binding site (RBS) of a target mRNA leading to translational repression (Bouvier et al., 2008; Jagodnik et al., 2017). Bacterial regulatory ncRNAs have notable roles in regulating several biological processes including the modulation of the bacterial cell wall (recently reviewed in Mediati et al., 2021), carbon metabolism (reviewed in Durica-Mitic et al., 2018) and virulence (reviewed in Sy et al., 2021). In *S. aureus*, the non-coding small RNA (sRNA) SprD, that is expressed from a pathogenicity island, was shown to repress the immune-evasion protein Sbi and is required for infection in a murine sepsis model (Chabelskaya et al., 2010). More recently, the sRNA RsaX28 (Ssr42) was implicated in the murine model of skin and soft tissue infection and regulates the expression of multiple virulence factors including the α and γ haemolysins, and capsule protein Cap5a through indirect regulation of Rsp (Das et al., 2016; Morrison et al., 2012), and through direct interactions with the δ haemolysin and enterotoxin I transcripts (*hld* and *sei*) (McKellar et al., 2022).

The untranslated regions (UTRs) of bacterial mRNAs can act as regulatory elements by base-pairing with target mRNAs, affecting translation and transcript stability. In our previous work, we demonstrated that the unusually long 3’ UTR of the *vigR* mRNA mediates vancomycin tolerance by upregulation of the cell wall lytic transglycosylase IsaA (Mediati et al., 2022). Some 3’ UTRs of mRNAs have been found to expand through insertion of sequence repeats including *Alu* elements in eukaryotes (Mayr, 2017) and IS elements in bacteria (Menendez-Gil et al., 2020). In *S. aureus*, the genome contains *<u>St</u>aphylococcus <u>a</u>ureus* repeat insertions (STAR repeats) that are short, repetitive motifs often separated by spacer sequences (Cramton et al., 2000). While the distribution of STAR repeats varies between closely related Staphylococci species, *S. aureus* isolates of the same evolutionary lineage (i.e., same multi-locus sequence-type) maintain a similar arrangement of STAR repeats (Purves et al., 2012). STAR repeats have been linked to pathogenesis (Purves et al., 2012), however their function, acquisition, and mechanism of propagation all remain unclear.

We have recently profiled the *in vivo* RNA interactome associated with the double-stranded RNA-specific endonuclease RNase III and mapped these interactions to genomic elements in the clinical MRSA isolate JKD6009 (McKellar et al., 2022; Mediati et al., 2022). Surprisingly, we found that the *vigR* 3’ UTR functions as a regulatory mRNA ‘hub’ required for glycopeptide tolerance (Mediati et al., 2022). In this study we have used MS2-affinity purification coupled with RNA sequencing (MAPS) to provide a more focussed snapshot of the RNA interaction partners of the *vigR* 3’ UTR. We find that *vigR* 3’ UTR interacts with mRNAs involved in carbon metabolism, amino acid biogenesis, cell wall biogenesis and virulence. We confirm a direct mRNA-mRNA interaction for the target mRNA *dapE*, a succinyl-diaminopimelate desuccinylase required for lysine and cell wall peptidoglycan synthesis. With our earlier finding that the *vigR* 3’ UTR up-regulates IsaA, the data suggests that *vigR* may play a broader role in controlling cell wall metabolism in *S. aureus*. Deletion of the *vigR* 3’ UTR (*vigR*^Δ3’UTR^) significantly increased the virulence of VISA in a wax moth larvae model of pathogenesis and we find that an *isaA* deletion is completely attenuated. Sequence analysis of *vigR* from a cross-section of *S. aureus* sequence types indicated that the *vigR* 3’ UTR is highly variable and has expanded through the acquisition of STAR repeats. We propose that expansion of the 3’ UTR may create a binding site for ribonucleases or RNA-binding proteins that functionalise the UTR. Our study has uncovered an extended regulatory network for the regulatory mRNA *vigR* and reveals a novel pathway of virulence regulation that is required for *S. aureus* infection.

## RESULTS

### The long 3’ UTR of *vigR* has expanded through STAR repeats in *S. aureus*

In our earlier work, we demonstrated that the long 3’ UTR of *vigR* is required for vancomycin tolerance in the VISA strain JKD6008 (Mediati et al., 2022). To understand if the *vigR* 3’ UTR is broadly conserved in *S. aureus* isolates we examined sequence variation within the *vigR* mRNA across 58 *S. aureus* genomes that represented a cross-section of sequence-types and clonal complexes. The 5’ and 3’ boundaries of *vigR* were previously defined in our dRNA-seq and Term-seq analyses (Mediati et al., 2022) and were used to extract *vigR* mRNA sequences with 5’ and 3’ UTRs. Both the 5’ UTR and coding sequence (CDS) of *vigR* are highly conserved between the 58 *S. aureus* genomes and did not vary in length except for the *vigR* coding sequence in strains JKD6008 and JKD6009 (423 nt c.f. 378 nt) (Figure 1A). In these isogenic strains a SNP has introduced a premature stop codon that truncates the VigR protein by 15 amino acids (Figure 1A). VigR is a hypothetical protein and it is not clear if the truncated VigR is functional, however these results suggest that *vigR* may be a pseudogene in JKD6008 and JKD6009, and may partly explain why we did not observe an antibiotic sensitivity phenotype for the *vigR*^ΔCDS^ deletion in our earlier analysis (Mediati et al., 2022).

**Figure 1.**
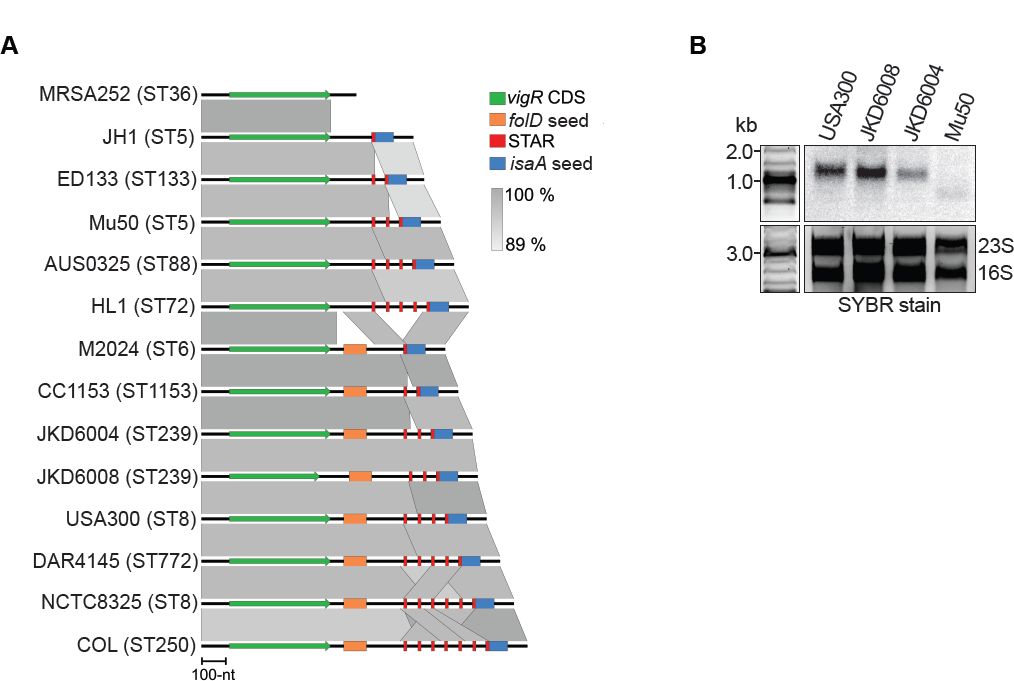
The *vigR* 3’ UTR varies in length between *S. aureus* isolates. **A**. Genomic alignment of the *vigR* transcript within 14 representative *S. aureus* strains extracted from dRNA-seq and Term-seq analyses. The VigR coding sequence (CDS) is represented in green and sequence expansion elements are indicated for the *folD* mRNA interaction seed (orange), *isaA* mRNA interaction seed (blue) and STAR repeat elements (red) (*right*) within the 3’ UTR. The degree of sequence conservation is indicated (*right*). **B**. Northern blot analysis of the *vigR* transcript. Total RNA was purified from *S. aureus* isolates indicated (*top*) and probed for the *vigR* CDS. SYBR Green stained 23S and 16S ribosomal RNAs are indicated below as loading controls.

In contrast to the 5’ UTR and CDS, the length of the *vigR* 3’ UTR varied from 102 – 819 nt across *S. aureus* genomes (**Supplementary Table 1**). To confirm that the *vigR* 3’ UTR varied between *S. aureus* isolates we performed Northern blot analysis on RNA extracted from strains USA300 (mRNA=1191 nt), JKD6008 (mRNA=1154 nt), JKD6004 (mRNA=1131 nt) and Mu50 (mRNA=999 nt) (Figure 1B and **Supplementary Figure 1**). In these strains we verified that the *vigR* mRNA transcript varied in length consistent with the variation predicted within the 3’ UTR from our sequence analysis (Figure 1A).

We next examined the *vigR* 3’ UTR to identify sequences that were responsible for expansion. Alignment and visualisation of 14 *vigR* mRNA sequences that represent the diversity of 3’ UTR lengths indicated that expansion had occurred in two regions (Figure 1A). The first region encompasses 162 nt at genomic positions 1,850,825 – 1,850,986 nt (using strain COL as a reference) (compare strains HL1 and M2023, insertion indicated in orange, Figure 1A). This site has introduced a mRNA interaction seed region with complementarity to the target mRNA *folD* (Mediati et al., 2022). The second region spans genomic positions 1,850,277 – 1,850,705 (in strain COL) and contains the predicted seed for the target mRNA *isaA* (indicated in blue, Figure 1A). This entire region also contains a repeated sequence that has expanded between *S. aureus* isolates (red, Figure 1A). Examination of the repeated sequences indicated the presence of a previously described STAR sequence repeat element with the consensus 5’ – TNTGTTGNGGCCCN – 3’ (Cramton et al., 2000). Among the genomes analysed, the *vigR* 3’ UTR contained 0 – 7 STAR repeats separated by ∼40 nt “spacer” sequences.

The spacer sequences between consecutive STARs are reported to be poorly conserved compared to the STAR motif (Purves et al., 2012). To better characterise the spacer-STAR sequences in *S. aureus* strain JKD6008, we used GLAM2 and GLAM2SCAN (Frith et al., 2008) to identify 101 spacer-STAR repeats throughout the entire JKD6008 genome and assembled a consensus sequence motif (Figure 2A and **Supplementary Table 2**). In line with previous studies, we find that the STAR motif is well conserved and our analysis extends the 5’ end of the STAR consensus by 4 nt to 5’ – <u>TCTN</u>TGTTGNGGCCCN – 3’. In addition, we find that the 12 nt at the 5’ end of the spacer is also well conserved among the 101 spacer-STAR sequences (Figure 2A).

**Figure 2.**
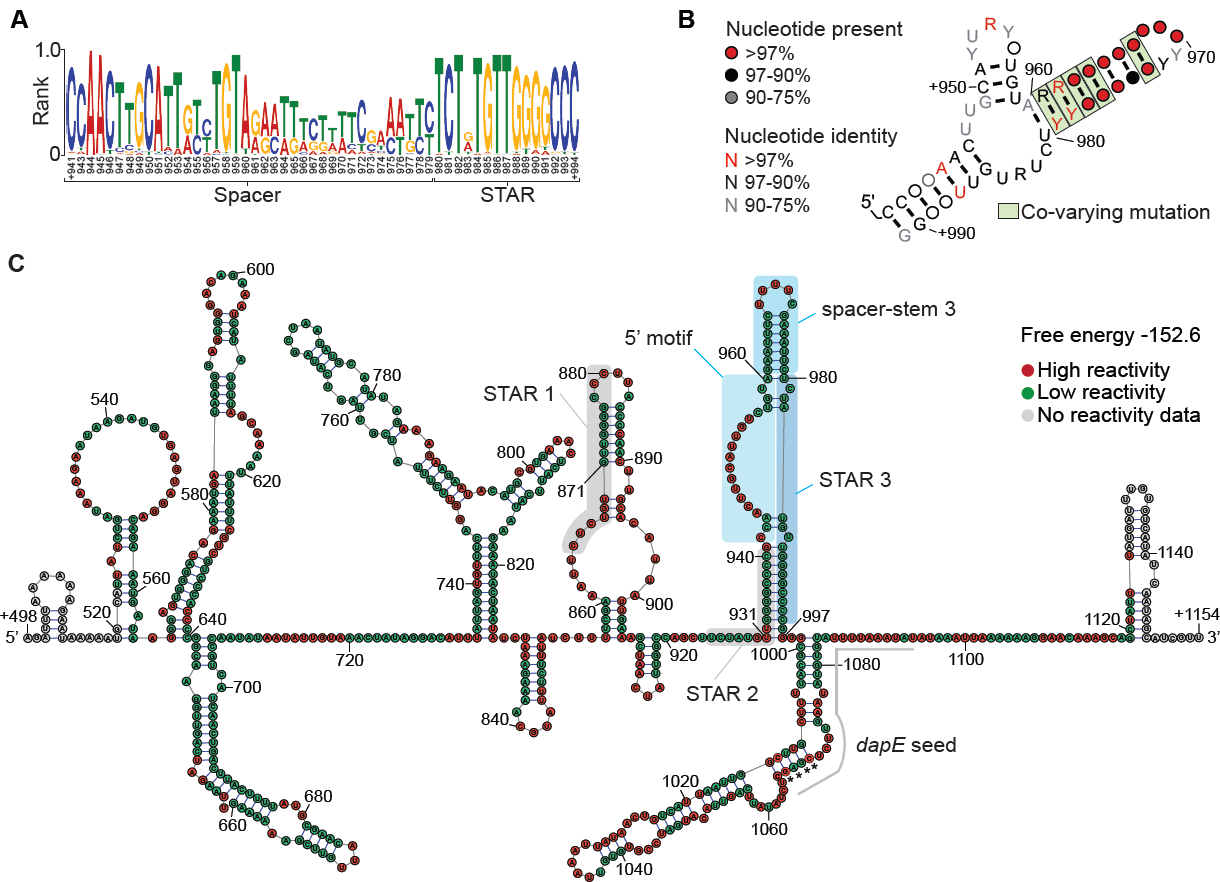
Spacer-STAR sequence repeats encode a conserved RNA structure. **A**. Consensus sequence motif of 101 spacer-STAR repeats identified within VISA isolate JKD6008 determined using GLAM2 software. The numbering of positions in the motifs are based on the numbering of spacer-STAR 3 of *vigR* (see panel 2C). **B**. Consensus RNA structural motif of spacer-STAR repeats using CMFinder software identifies co-varying nucleotides. Statistically significant covariation (*green*) was determined using R-scape software. The numbering of positions in the motifs are based on the numbering of spacer-STAR 3 of *vigR* (see panel 2C). Probability of nucleotide presence and identities are indicated (*left*). **C**. The *in vitro* secondary structure of the *vigR* 3’ UTR from VISA isolate JKD6008. Benzyl cyanide and lead acetate were used to modify the RNA backbone and nucleotide reactivity (*right*) was determined by separation on TBE-urea gels. The nucleotide positions are relative to the transcription start site of *vigR*. STAR motif 1 and 2 are indicted by grey boxes. Spacer-STAR repeat 3 is indicated by blue boxes. Each section (5’ motif, spacer-stem, and STAR motif) are indicated for spacer-STAR 3. The nucleotides predicted to interact with *dapE* mRNA are indicated by the grey line. Asterisks indicated the positions where the *isaA* and *dapE* interaction sites overlap.

### STAR spacer sequences contain a conserved RNA structure

Given the expansion of spacer-STAR repeats in the *vigR* 3’ UTR we next asked if spacer-STAR loci are transcribed in other genomic contexts. Spacer-STAR sequences were mapped to the *S. aureus* transcriptome, and we found that 24 were within 3’ UTRs, 10 within 5’ UTRs, and 17 within predicted sRNAs (defined by SRD, Sassi et al., 2015) (**Supplementary Figure 2A**). The remaining 50/101 were within intergenic regions but not within our experimentally defined transcriptome boundaries (Mediati et al., 2022). These results indicate that many spacer-STARs are inserted into UTRs and non-coding sRNAs suggesting that spacer-STAR sequences may encode a functional RNA.

To determine if spacer-STARs encode conserved RNA structure, we used CMFinder (Yao et al., 2006) to identify co-varying nucleotides indicative of conserved structure within our 101 spacer-STAR sequences in *S. aureus* strain JKD6008. Consensus RNA structures and sequences were analysed for statistically significant covariation using R-scape (Rivas et al., 2020) and visualised using R2R (Weinberg and Breaker, 2011). We identified 5 statistically significant co-varying bases positioned within a single stem-loop of the spacer-STAR (Figure 2B and **Supplementary Table 3**). This conserved RNA stem-loop structure is positioned from +960 – 980 nt of the *vigR* 3’ UTR in our GLAM2 motif (Figure 2A) and corresponds to the poorly conserved spacer sequence. These data indicate that while there is low sequence conservation in the spacer, an 8 base-pair long RNA stem-loop structure is conserved. While not statistically significant (likely due to high sequence conservation), the conserved 5’ spacer and 3’ STAR sequences are predicted to form an RNA duplex at the base of the structure (Figure 2B).

Collectively, these data indicate that the *vigR* 3’ UTR has expanded within *S. aureus* genomes through insertion of a 162 nt sequence and spacer-STAR repeats. While the sequence of the STAR repeats and 5’ end of the spacer are conserved, positions +960 – 980 nt of the spacer encodes an 8 base-pair long RNA stem-loop with variable sequence suggesting that the RNA structure – rather than the sequence - of the spacer is functionally important.

### Spacer-STAR repeats are structured *in vitro*

To confirm that the spacer-STAR sequence forms a conserved RNA structure, we used benzyl cyanide and lead acetate to probe the *in vitro* secondary structure of the *vigR* 3’ UTR from *S. aureus* strain JKD6008 that contains 3 spacer-STAR repeats (Figure 2C and **Supplementary Figure 2B**). Local nucleotide reactivity and flexibility was analysed using RNAstructure software (Reuter and Mathews, 2010) to predict secondary structure within the *vigR* 3’ UTR. We find that the *vigR* 3’ UTR is highly structured and that the third spacer-STAR repeat (STAR 3) forms the stem-loop structure predicted by sequence co-variation (Figure 2B), albeit with base-pairing between the STAR motif of repeat 2 and 3, rather than the conserved 5’ motif and STAR 3 (shaded blue in Figure 2C). The spacer-stem of STAR repeat 2 is partially retained (+906-919, Figure 2B), but appears to have been lost from STAR repeat 1 in our structure prediction. Overall our *in vitro* structure probing data of *vigR* 3’ UTR support formation of at least one spacer-STAR stem-loop structure predicted by sequence co-variation.

### *vigR* 3’ UTR represses the *dapE* mRNA *in vivo*

The *vigR* 3’ UTR was previously shown to interact with the *folD* and *isaA* mRNAs (Mediati et al., 2022), the latter encodes a lytic transglycosylase that cleaves the β-1,4-glycosidic bonding between the *N*-acetylmuramic acid (MurNAc)-*N*-acetylglucosamine (GlcNAc) residues of cell wall peptidoglycan. While deletion of *isaA* reduced cell wall thickness and conferred sensitivity to the glycopeptide antibiotic teicoplanin, the *isaA* mutation does not confer the same vancomycin sensitivity seen in the *vigR* 3’ UTR deletion (Mediati et al., 2022), suggesting that *vigR* may have additional targets in *S. aureus* strain JKD6008. To identify interaction partners for the *vigR* 3’ UTR we used MS2-affinity purification and sequencing (MAPS) (Lalaouna et al., 2015; Said et al., 2009). The MS2 RNA aptamer was fused to the 5’ end of the *vigR* 3’ UTR and placed under the control of the P*_xyl/tet_* promoter of pRAB11 (Helle et al., 2011). Inducible transcription of MS2-*vigR* 3’ UTR was confirmed by Northern blot, and we find that 15 min of induction with anhydrotetracycline (ATc) leads to strong accumulation of the MS2 fusion (**Supplementary Figure 3**). The MS2 fusion was induced in *S. aureus* strain JKD6008 grown in BHI media to an OD_600nm_ 3.0. (mid-log growth phase). Cells were pelleted and lysed before loading onto an amylose column loaded with MS2 protein-His-MBP fusion to pull-down the MS2 aptamer. After washing, bound RNAs were eluted with maltose and precipitated for library preparation and sequencing (Lalaouna et al., 2015; Mercier et al., 2021). Peaks were called within the sequencing datasets using blockbuster (Langenberger et al., 2009) and CRAC software (Webb et al., 2014). DESeq2 was used to identify 81 statistically significant peaks enriched in the duplicate MS2-*vigR* 3’ UTR samples compared to MS2-only controls (*p*<0.01) (**Supplementary Table 4**). Clusters of orthologous group (COG) analyses for statistically significant transcripts enriched in MS2-*vigR* 3’ UTR MAPS indicated that functional classifications associated with “Carbohydrate transport and metabolism”, “Amino acid transport and metabolism” and “Cell wall, membrane and envelope biogenesis” were enriched (adjusted *p*<0.05, Figure 3A). To identify interactions that affect the abundance of target RNAs, we correlated our MAPS enrichment data with RNA-seq differential expression data from JKD6008 *vigR*^Δ3’UTR^ (Mediati et al., 2022). A total of 22 mRNA transcripts were enriched >2-fold by MAPS and had >2-fold increased expression in the 3’ UTR deletion strain (Figure 3B, red dotted line), suggesting that *vigR* 3’ UTR may repress these mRNAs through a direct RNA-RNA interaction.

**Figure 3.**
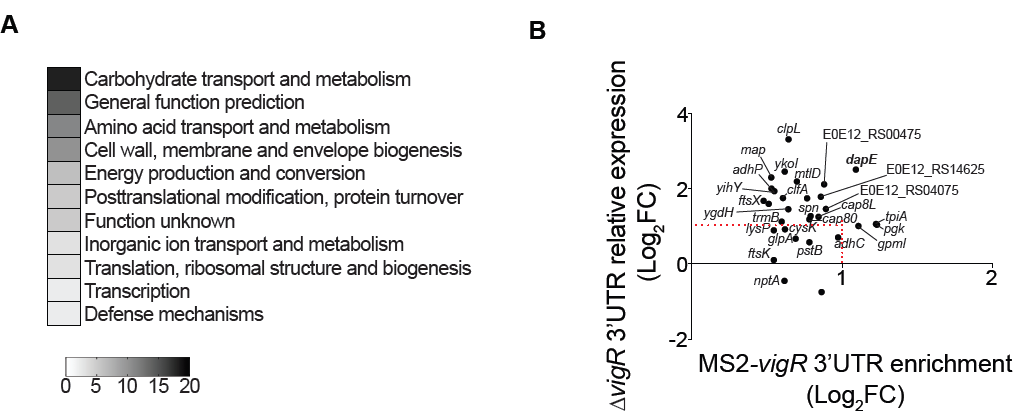
MAPS identifies the extended regulon of *vigR* 3’ UTR in VISA JKD6008. **A**. Clusters of orthologous groups (COGs) detailed for the 81 statistically significant enriched transcripts found in duplicate MS2-*vigR* 3’ UTR MAPS experiments (*p*<0.01). The most abundant COGs are listed in numerical order. **B**. Correlation of enriched transcripts from MAPS (*bottom*) with dysregulated transcripts from RNA-seq of JKD6008 *vigR*^Δ3’UTR^ (*left*). The red dotted line indicates those transcripts with a log_2_FC ≥1 in both MAPS and RNA-seq.

We used electrophoretic mobility shift assays (EMSAs) to verify a direct RNA-RNA interaction *in vitro*. The mRNAs *dapE*, *spn* and *hysA* that represent different levels of enrichment by MAPS an RNA-seq were *in vitro* transcribed from JKD6008 and incubated with radiolabelled *vigR* 3’ UTR before separation on native 4% TBE PAGE gels. Only the *dapE* mRNA (SAA6008_RS11085) was gel shifted by *vigR* 3’ UTR and we find that the complex formed between these long RNAs (657-nt and 1,295-nt, respectively) does not migrate out of the well (Figure 4A, EMSA for *spn* and *hysA* mRNAs in **Supplementary Figure 4A**). To identify the interaction site, we divided the *dapE* mRNA into 3 sub-fragments and repeated the EMSA (**Supplementary Figure 4B**). The *vigR* 3’ UTR was able to gel shift *dapE* Frag-B (462 nt length) encompassing genomic positions 2,162,029 to 2,162,473 nt (Figure 4B **and Supplementary Figure 4C**). To further narrow down the interaction site, antisense competitor oligonucleotides (termed 1-4) were tiled across the interaction site between Frag-B and *vigR* 3’UTR (Figure 4C-D **and Supplementary Figure 4D**). The 31-nt antisense oligo 4 was able to compete away *dapE* Frag-B from *vigR* 3’ UTR (Figure 4C). This site contains a predicted 28 base-pair interaction between *dapE* and *vigR* 3’ UTR, with 22-nt of complementarity (Figure 4E). Notably, the *dapE* seed sequence (+1066–1094 nts) partially overlaps the *isaA* mRNA seed region at +984 – 1069 (Figure 2C)(Mediati et al., 2022). This data suggests that this region within the *vigR* 3’ UTR can base-pair with multiple RNA targets.

**Figure 4.**
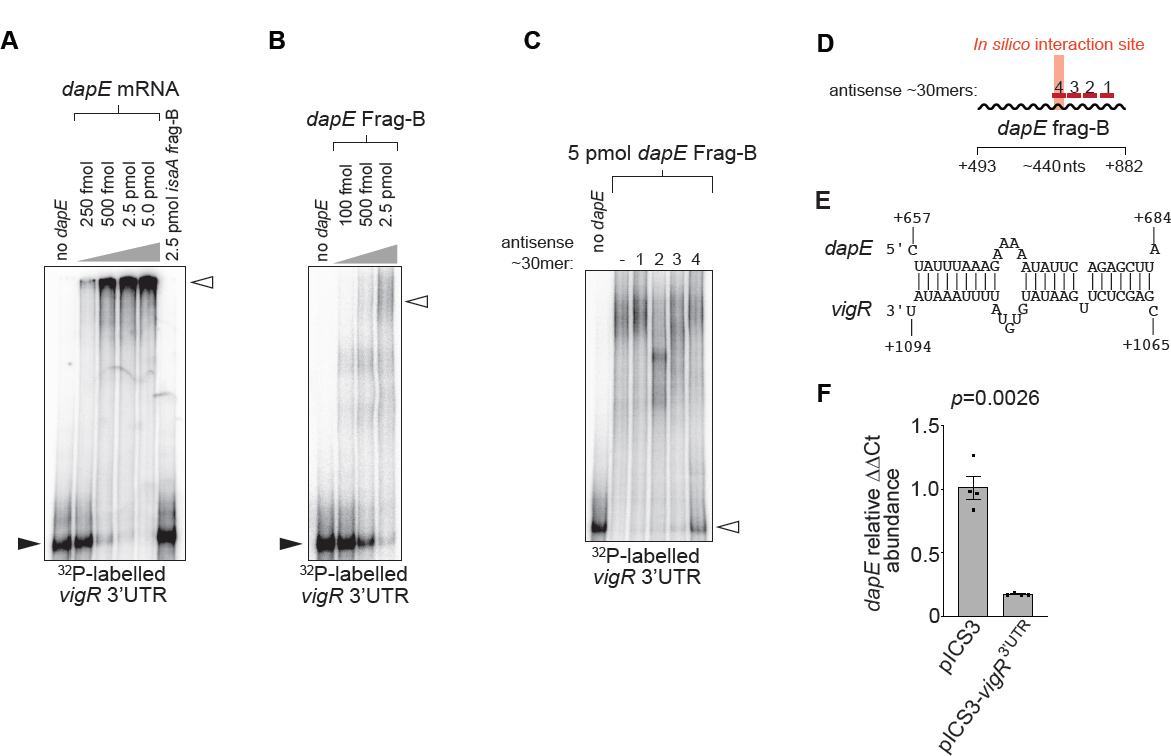
The succinyl-diaminopimelate desuccinylase *dapE* is regulated by *vigR* 3’ UTR. **A**. EMSA analysis of the RNA-RNA interaction between *vigR* 3’ UTR and *dapE* mRNA. A total of 50 fM of radiolabelled *vigR* 3’ UTR (*bottom*) was titrated against increasing concentrations of the *dapE* mRNA (*top*). The *isaA* fragment B (frag-B) RNA was titrated against 50 fM of radiolabelled *vigR* 3’ UTR as a known negative control. **B**. The *dapE* mRNA was synthesised as sub-fragments (∼400-nt in length) and used for EMSA analysis. *dapE* fragment B RNA and concentrations are indicated (*top*). Black arrowheads indicate migration of free, radiolabelled *vigR* 3’ UTR and open arrowheads indicate slow migrating *vigR*-*dapE* duplexes. **C**. EMSA analysis of interactions between *vigR* 3’ UTR and *dapE* frag-B (0 or 2.5 pmol). Antisense competitor oligonucleotides 1-4 (*top*) were spiked in at 500x excess concentration. **D**. The antisense oligonucleotide competitors 1-4 used for EMSA analysis are indicated relative to the *dapE* frag-B RNA. The start and end positions of *dapE* frag-B are indicated representative of the *dapE* transcription start site (+1 site). The predicted *in silico* interaction site is indicated in red (*top*). **E**. The predicted interaction seed between the *vigR*-*dapE* RNA species. The start and end positions of the RNA-RNA duplex are indicated representative of the mRNA transcription start sites. **F**. Histogram of quantitative RT-PCR to quantify *dapE* chromosomal abundance (relative to *gapA*) in the pICS3 and pICS3::*vigR*^3’UTR^ constructs (*top*). Error bars represent standard error of the mean (SEM). *p*=0.0026, *n*=4.

To verify that the interaction between *vigR* 3’ UTR and *dapE* mRNA is functional, we overexpressed *vigR* 3’ UTR from the P*_tufA_* promoter of pICS3 (Ivain et al., 2017) and assessed *dapE* mRNA abundance using qRT-PCR (Figure 4F). Consistent with our earlier RNA-seq analysis, we find that *dapE* is 6.1-fold repressed by the *vigR* 3’ UTR (*p*=0.0026, *n*=4, Figure 4F).

Collectively, these data indicate that the *vigR* 3’ UTR represses *dapE* through a direct base-pair interaction with the coding sequence of the mRNA. DapE encodes succinyl-diaminopimelate desuccinylase that is required for lysine and peptidoglycan synthesis (Born and Blanchard, 1999) and our results indicate that in addition to activation of the cell wall lytic transglycoslase *isaA*, *vigR* represses *dapE* that contributes to cell wall biosynthesis.

### The *vigR* 3 UTR is required for vancomycin tolerance and virulence in a wax moth larvae model of infection

Intermediate-vancomycin resistance in *S. aureus* isolates has been correlated with a decrease in virulence in both murine bacteraemia and wax moth larvae models of infection (Cameron et al., 2017; Jin et al., 2020). We next asked if the decreased vancomycin tolerance of our *vigR*^Δ3’UTR^ strain also results in increased virulence in a wax moth larvae model of infection. We infected larvae (*n*=20) with 10 µL of 10^7^ CFU/ml of VISA strain JKD6008 (isogenic parent), the *vigR*^Δ3’UTR^ strain, and marker rescue strain *vigR*^Δ3’UTR^-repair (*vigR*^Δ3’UTR::3’UTR^) where the wild-type 3’ UTR sequence has been restored. We also included the vancomycin-sensitive (VSSA) strain JKD6009 that is the parent strain of JKD6008 (Howden et al., 2010). Infected larvae were monitored for 6 days for melanisation and death. Consistent with earlier studies (Cameron et al., 2017; Jin et al., 2020), we find that the VSSA strain JKD6009 is significantly more virulent than the VISA derivative JKD6008 (*p*=0.0001, Figure 5A). Deletion of the *vigR* 3’ UTR significantly increased the virulence of JKD6008 (*p*=0.031), and wild-type virulence was restored in the *vigR*^Δ3’UTR^-repaired strain (Figure 5A). This result indicates that the 3’ UTR of *vigR* contributes to both the reduced virulence and vancomycin intermediate resistance of VISA strain JKD6008.

**Figure 5.**
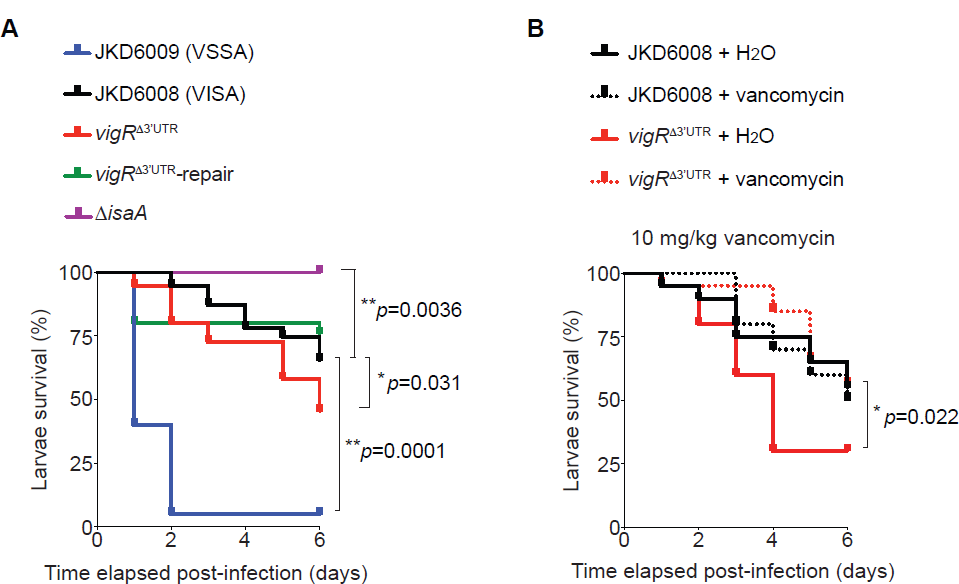
The *vigR* 3’ UTR and *isaA* mRNA are required for pathogenesis. **A**. Kaplan-Meier survival plot of *Galleria mellonella* larvae infected with 10^7^ CFU of *S. aureus* constructs (*top)* over the course of 6 days. Plots show an average of 4 independent replicates with 5 larvae per replicate (*n*=20). Significant differences between survival curves were determined by Log-rank test at \**p*<0.05 and \*\**p*<0.005. **B**. The *vigR* 3’ UTR contributes to vancomycin tolerance during infection. Larvae were infected with 10^7^ CFU of JKD6008 and *vigR*^Δ3’UTR^ strains and challenged with either 10 mg/kg of vancomycin or H_2_O over the course of 6 days. Plots show an average of 2 independent replicates with 10 larvae per replicate (*n*=20). Significant differences between curves were determined by Log-rank test at \**p*<0.05.

To understand whether the *vigR* 3’ UTR contributes to vancomycin tolerance during infection, we treated larvae with 10mg/kg of vancomycin directly after injection with VISA JKD6008 or the *vigR*^Δ3’UTR^ strain (Figure 5B). Consistent with *in vitro* results, treatment with vancomycin did not significantly affect the virulence of the vancomycin-intermediate strain JKD6008. However, vancomycin treatment significantly reduced killing in the *vigR*^Δ3’UTR^ strain and reduced virulence to wild-type levels (*p*=0.022, Figure 5B). The addition of 10mg/kg of vancomycin delayed complete killing of larvae infected with VSSA isolate JKD6009 by 1-day and did not significantly affect pathogenesis (**Supplementary Figure 5**). These results indicate that the 3’ UTR of *vigR* is required for intermediate vancomycin resistance in VISA during infection.

We had previously shown that the 3’ UTR of *vigR* up-regulates the cell wall lytic transglycosylase *isaA* that contributes to cell wall thickening in VISA JKD6008 (Mediati et al., 2022). We additionally infected larvae with the JKD6008 Δ*isaA* strain to determine if *vigR* 3’ UTR regulation of *isaA* contributes to the virulence phenotype. In contrast to the *vigR* 3’ UTR, deletion of *isaA* completely attenuated JKD6008 (*p*=0.0036, Figure 5A). Our results demonstrate that while deletion of *isaA* reduces cell wall thickness comparable to the virulent VSSA strain JKD6009, virulence is not restored to VSSA levels in the Δ*isaA* background. Our data indicate that the cell wall lytic transglycosylase IsaA plays a critical role in *S. aureus* infection.

## DISCUSSION

In earlier work, the regulatory 3’ UTR of *vigR* mRNA was found to control vancomycin tolerance and upregulate the lytic transglycosylase *isaA* (Mediati et al., 2022). Using MS2-affinity purification and RNA sequencing, we demonstrate that the *vigR* 3’ UTR also represses *dapE*, a succinyl-diaminopimelate desuccinylase that is required for lysine and peptidoglycan synthesis (Gillner et al., 2013). Our results suggest that the *vigR* 3’ UTR may play a broader role in controlling cell wall metabolism in *S. aureus* and we demonstrate that the *vigR* 3’ UTR also contributes to the attenuated virulence of the vancomycin-intermediate isolate. Surprisingly, we find that *isaA* is not required for growth *in vitro*, but the Δ*isaA* strain is completely attenuated in a wax moth larvae model of pathogenesis.

Sequence analysis of *vigR* indicated that the 5’ UTR and CDS are highly conserved. In contrast, the 3’ UTR is highly variable and appears to expand through the acquisition of STAR repeats (Cramton et al., 2000). Here we have extended the reported 14-nt STAR repeat sequence to include a conserved 5’ 4-nt <u>TCTN</u> and we find that while the ∼40 nt variable spacers do not have conserved sequence, they form an evolutionary conserved RNA stem-loop structure suggesting that the structure of the spacer region is functional. Our *in vitro* structure probing data supports the formation of an RNA stem in the spacer region.

It is not yet clear how the spacer-STAR elements influence the function of the *vigR* 3’ UTR, but it is notable that repeat elements have previously been linked to mRNA stability in bacteria and eukaryotes (Chan et al., 2022; De Gregorio et al., 2002; De Gregorio et al., 2005; De Gregorio et al., 2006; Knutsen et al., 2006; Maquat, 2020; Menendez-Gil et al., 2020). Sequence repeats termed SINE elements (notably *Alu* elements in humans) are known to modulate RNA-RNA and RNA-protein interactions when inserted into eukaryotic 3’ UTRs (Chan et al., 2022; Maquat, 2020). *Alu* elements in 3’ UTRs are reported to provide interaction sites for the dsRNA ribonuclease Staufen (Gong and Maquat, 2011; Lucas et al., 2018). In a mechanism that may functionally parallel our observations with *vigR*, 3’ UTR-encoded *Alu* repeats facilitate interactions with *Alu*-encoding long non-coding RNAs (lncRNAs) (Gong and Maquat, 2011). Imperfect base-pairing of the 3’ UTR and lncRNA recruits Staufen, triggering Staufen-mediated decay and repression of the mRNA target. The *isaA* interacting nucleotides in *vigR* partially overlap STAR repeat 3 of *vigR* indicating that this interaction is partly driven by acquisition of the STAR repeat. By analogy, the imperfect base-pairing between the *vigR* STAR repeats and mRNA targets may create a binding site for ribonucleases or recruit RNA binding proteins.

In our previous work we postulated that the *vigR* 3’ UTR interactions with *isaA* and *folD* mRNAs may occlude an RNase cleavage site to stabilise the transcripts. This is in line with previous work showing that the regulatory mRNA-mRNA interactions between *hyl*-*prsA* and *irvA*-*gbpC* stabilise their targets by occluding interactions with RNase J1 (Ignatov et al., 2020; Liu et al., 2015). Here we find that *vigR* represses *dapE*, suggesting a mechanism where the *vigR*-*dapE* interaction may create an RNase targeting site, although we have not identified the RNase responsible. To our knowledge, this is the first repressive regulatory mRNA-mRNA interaction identified in bacteria and indicates that these interactions can have both activating and repressing regulatory outcomes.

By intersecting RNA-seq and MAPS data we have uncovered a functional interaction between *dapE* mRNA and the *vigR* 3’ UTR. DapE is a succinyl-diaminopimelate desuccinylase that is required for lysine and peptidoglycan synthesis (Gillner et al., 2013). Repression of *dapE* and activation of *isaA* expression (a cell wall autolysin) suggests that *vigR* upregulation coordinates a reduction in cell wall peptidoglycan crosslinking. Lysine is required for transpeptidation of peptidoglycan in *S. aureus*, and IsaA cleaves the glycosidic bonds between the MurNAc and GlcNAc sugars within peptidoglycan strands. This coordinated reduction in peptidoglycan crosslinking would be consistent with the reduced cell wall crosslinking that is observed in VISA strains (Howden et al., 2010).

Cell wall thickening and reduced crosslinking is thought to contribute to vancomycin tolerance in VISA strains however, these strains are generally less virulent in wax moth and mouse models of infection (Cameron et al., 2017; Jin et al., 2020). We have confirmed this phenotype for VISA JKD6008 and found that upregulation of *vigR* in the VISA strain partially contributes to both vancomycin tolerance *in vivo*, and to the reduced virulence phenotype. To our surprise, deletion of the lytic transglycosylase *isaA* completely attenuated virulence in the wax moth model. Previous studies have demonstrated that passive immunisation with IsaA-targeting IgG (Lorenz et al. 2011) can reduce mortality in a mouse model of infection, indicating that IsaA is presented on the surface of the cell during infection, and our data suggest that IsaA may also contribute to virulence in VISA strain JKD6008, although the mechanism remains unclear. Cell wall metabolism appears to change during infections and the ability to crosslink and cleave peptidoglycan plays an important role in virulence (Sutton et al., 2021). The glucosaminidase SagB (that cleaves peptidoglycan) is required for virulence in a mouse model of infection although the precise mechanism is also unknown (Sutton et al., 2021). The cell wall autolysin LytM is required for release of Protein A in *S. aureus*, linking cell wall hydrolysis to virulence (Becker et al., 2014) and suggesting a potential mechanism for Δ*isaA* attenuation - through release of virulence factors at the cell surface.

Collectively, our data indicates that the long 3’ UTR of *vigR* has been functionalised by the acquisition of STAR sequence repeats that encode structured RNA. The JKD6008 genome encodes at least 101 STAR repeats and based on our earlier mapping of transcript boundaries (Mediati et al., 2022), we predict that 51 of these are transcribed (data not shown). Within the context of the *vigR* 3’ UTR, the STAR repeat facilitates interactions with both *dapE* and *isaA* mRNAs that is predicted to reduce cell wall peptidoglycan crosslinking. The *vigR* 3’ UTR reduces virulence in a wax moth model of infection, consistent with the VISA phenotype – but this does not appear to be dependent on up-regulation of *isaA* that is required for larvae killing. Our results support a broader role for *vigR* 3’ UTR in regulation of cell wall metabolism that contributes to both vancomycin tolerance and reduced virulence in VISA.

## METHODS

### Bacterial strains and general culture conditions

The bacterial strains, plasmids and oligonucleotides used in this study are listed in **Supplementary Table 5**. *S. aureus* strains RN4220, USA300, Mu50, JKD6004 and the JKD6009/JKD6008 (VSSA/VISA) pair were routinely cultured at 37°C on solid or in liquid brain heart infusion (BHI, Merck) or Mueller-Hinton (MH, Merck) media. Antibiotics were used in this study to select for plasmids in *S. aureus* at 15 μg/mL chloramphenicol, unless otherwise specified. *E. coli* DH5a and IM08B strains were cultured at 37°C on solid or in liquid Luria-Bertani (LB) media. Antibiotics were used to select for plasmids in *E. coli* at 100 μg/mL ampicillin or 15 μg/mL chloramphenicol, unless otherwise specified. All bacterial strains were stored at −80°C as stationary phase cultures with 16% (v/v) glycerol.

### Strain modifications and plasmids

*S. aureus* MS2-affinity tagged constructs were constructed in the anhydrotetracycline (ATc) inducible P_xyl/tet_ pRAB11 vector system (Helle et al., 2011). The *vigR* 3’ UTR sequence was amplified from JKD6009 using Phusion Hot Start Polymerase (NEB) with primers incorporating the MS2 aptamer sequence (fused to 5’ end of *vigR* 3’ UTR) and the *rrn1* T7 terminator (**Supplementary Table 5**). The MS2-*vigR* 3’ UTR and MS2 products were cloned into pRAB11 at the KpnI and EcoRI sites using 10 U of T4 DNA ligase (Thermo) and transformed into chemically-competent *E. coli* DH5α. Antibiotics were used to select for pRAB11 in *E. coli* at 100 μg/mL ampicillin. Constructs were confirmed by Sanger sequencing, transformed into electrocompetent *E. coli* IM08B and then transformed into electrocompetent *S. aureus* JKD6009 *vigR*^Δ3’UTR^ (Mediati et al., 2022). Antibiotics were used to select for pRAB11 in *S. aureus* at 15 μg/mL chloramphenicol. Construction of the *S. aureus* JKD6009 *vigR*^Δ13’UTR^, JKD6009 pICS3::*vigR,* JKD6008 *vigR*^Δ13’UTR^, JKD6008 *vigR*^Δ13’UTR^-repair, JKD6008 Δ*isaA* strains were described previously in Mediati et al., 2022.

### *vigR* conservation analysis

The mRNA transcriptional boundary of *vigR* in *S. aureus* strain JKD6009 was determined from our previous work using dRNA-seq, Term-seq and Northern blot (Mediati et al., 2022). These boundary nucleotides were used to define and extract the *vigR* mRNA sequence from 58 genomes of *S. aureus* isolates. The 5’ UTR, CDS and 3’ UTR sequence length, number of STAR repeats, and presence or absence of the *folD* or *isaA* mRNA interaction seed region in each isolate were determined individually and used to construct **Supplementary Table 1**. From these 58 genomes, 14 representative strains were selected that demonstrated *vigR* transcript variation. The GenBank and blastn.out files from these 14 representative strains were used as input into Easyfig (Sullivan et al., 2011) to generate Figure 1A.

### Northern blot

Total RNA was purified using the GTC-phenol:chloroform extraction method as performed previously for *S. aureus* (Mediati et al., 2022). At least 5 μg of RNA was treated with a 5:1 ratio of glyoxal denaturation mixture for 1 h at 55°C. Denatured RNA was resolved on a 1% BPTE-agarose gel containing SYBR Green (Thermo) and run for ∼1 h at 100 V in 1x BPTE buffer. Intact 23S and 16S ribosomal RNA was confirmed on a Bio-Rad Chemi-doc and washed consecutively in 200 mL of 75 mM NaOH, 200 mL of neutralizing solution (1.5 M NaCl and 500 mM Tris-HCI, pH 7.5) and 200 mL of SSC buffer (3 M NaCl and 300 mM sodium citrate, pH 7.0) for 20 min each. RNA was capillary transferred onto a Hybond-N+ nylon membrane (GE Healthcare) and UV-crosslinked in a Stratagene Auto-Crosslinker with 1200 mJ dosage of UV-C. The membrane was equilibrated in Ambion ULTRAhyb hybridization buffer (Thermo) for 1 h at 42°C and then incubated with 10 pMol of 20 μCi γ^32^P-ATP-labelled oligonucleotide probe (**Supplementary Table 5**) for 16 h at 42°C. Membranes were washed three times in 2x sodium chloride sodium phosphate EDTA (SSPE) buffer with the addition of 0.1% SDS for 20 min at 42°C. The blot was imaged using a BAS-MP 2040 phosphorscreen on a FLA9500 Typhoon (GE Healthcare). ImageJ software (Schindelin et al., 2012) was used to align the SYBR stained gel and membrane, and used to construct Figure 1B.

### *In silico* RNA structure prediction

The STAR sequence repeats, as defined previously by (Purves et al., 2012) (5’ – TNTGTTGNGGCCCN), and the upstream 50 nt within JKD60008 were extrapolated and used as input into the GLAM2 software (Frith et al., 2008) to generate the consensus Spacer-STAR sequence motif in Figure 2A. GLAM2 was used with the gapless Gibbs sampling parameter and accommodates short sequence gaps. This consensus Spacer-STAR motif was inserted into GLAM2SCAN (Frith et al., 2008) and used to identify related sequences within the JKD6008 genome. The transcriptional boundaries as identified previously using dRNA-seq and Term-seq (Mediati et al., 2022) were then used to determine the genomic features (e.g., UTRs, CDS or sRNA) that each Spacer-STAR sequence is positioned in. These 101 Spacer-STAR sequences defined by GLAM2SCAN were then used as input into CMfinder software (Yao et al., 2006) and used to construct the *in silico* consensus RNA secondary structure model in Figure 2B using the covariance model expectation maximisation algorithm. The R-scape software program (Rivas et al., 2020) was used to assess statistical significance of the co-varying base pairs.

### *In vitro* structure of *vigR* 3’ UTR

Purified *vigR* 3’ UTR amplified from JKD6008 was *in vitro* transcribed (IVT) using HiScribe T7 RNA polymerase (NEB). RNA products were DNase I treated (NEB) for 30 mins at 37°C, phenol-chloroform extracted, ethanol precipitated, and then separated on a 4% polyacrylamide TBE-6M urea gel. Products were excised, crushed, and incubated in 500 μL gel elution buffer (10 mM magnesium acetate, 0.5 M ammonium acetate, 1 mM EDTA) with gentle rotation for 16 h at 4°C. RNA was extracted from the eluate using phenol-chloroform and ethanol precipitation. Approx. 5 pMol of purified *vigR* 3’ UTR RNA was renatured by heating to 90°C for 2 min, placed on ice for 2 min, and then incubated in folding buffer (300 mM HEPES (pH 8.0), 20 mM MgCl_2_ and 300 mM NaCl) for 1 h at 37°C. RNA was then chemically modified with 10, 50 and 100 mM of benzoyl cyanide (Sigma) for 1 min at 20°C. RNA species were also modified with 10 mM lead acetate (Sigma) for 1 min at 25°C. As a no-modification control, DMSO (Sigma) was added to the RNA and incubated for 1 min at 20°C. RNA species were ethanol precipitated and reverse transcribed using SuperScript IV (Thermo) with purified 30 μCi ^32^P-ATP-labeled oligonucleotides spanning the entire *vigR* 3’ UTR (**Supplementary Figure 2B** and **Supplementary Table 5**). In parallel, single ddNTP (Roche) sequencing reactions were performed with identical 30 μCi ^32^P-ATP-labeled oligonucleotides and 2 pMol of RNA. The cDNA products were incubated with 200 mM NaOH at 80°C to hydrolyse template RNA and inactivate SuperScript IV enzyme. Products were separated on a 6% polyacrylamide TBE-6M urea gel for 100 min at a maximum of 50 W (**Supplementary Figure 2B**). Gels were then dried and visualised using a Fuji BAS-MP 2040 phosphorscreen and Typhoon FLA9500. Nucleotide reactivity was analysed and RNAstructure (Reuter and Mathews, 2010) was used to construct the secondary structure model in Figure 2C.

### MS2-affinity purification and RNA-sequencing (MAPS)

JKD6009 *vigR*^Δ13’UTR^ transformed with pRAB11::MS2-*vigR*^3’UTR^ and pRAB11::MS2 (MS2 tag only control) were grown in BHI media supplemented with 15 μg/mL chloramphenicol at 37°C with 180 rpm shaking to an OD_600nm_ 3.0. Constructs were then induced with 0.4 μM ATc and grown for a further 15 min at 37°C with shaking. Cultures were harvested by centrifugation at 4°C and crude extracts (5 μg) were probed for the MS2 aptamer sequence. MS2-affinity purifications were performed in biological duplicates and as previously described (Lalaouna et al., 2015; Mercier et al., 2021). RNA quality was assessed on a PicoRNA Bioanalyzer 2100 chip and underwent ribosomal RNA depletion using QIAseq FastSelect (Qiagen). Sequencing libraries were constructed using the NEBNext II directional RNA library kit for Illumina sequencing (NEB) and sequenced on a NextSeq2000 platform at the Epitranscriptomics and RNA-sequencing facility, Université de Lorraine-CNRS-INSERM (Nancy, France) generating 50 bp single-end reads.

### Analysis of enriched nucleotide peak data

MAPS data were analysed using the pipeline previously described for CRAC data analysis (Sy et al., 2018) using the ruffus pipeline CRAC_pipeline_SE.py that performs alignment and read counting steps (https://git.ecdf.ed.ac.uk/sgrannem/crac_pipelines). Binding sites were identified using blockbuster as described previously in Sy et al., 2018 and adapted from Holmqvist et al., 2016. Briefly, GTF format outputs from the ruffus CRAC pipeline were converted to BED format using pyGTF2bed.py (Webb et al., 2014). The experimental replicates were combined and sorted before peak calling. Peaks were defined using blockbuster with settings: - minBlockHeight 50 -distance 1. Peak intervals defined by blockbuster were used to calculate statistically enriched regions of the transcriptome. Read depth at peak intervals was calculated for each experimental and control replicate using HTSeq (Anders et al., 2015), and enriched peaks identified using DESeq2 (Love et al., 2014).

### RNA-RNA electrophoretic mobility shift assay (EMSA)

Full-length or sub-fragments of *dapE* (SAA6008_RS11085), *spn* (SAA6008_RS02260), *hysA* (SAA6008_RS12255) and *vigR* 3’ UTR from JKD6008 were *in vitro* transcribed (IVT) using HiScribe T7 RNA polymerase (NEB). IVT products were RQ1 DNase treated (Promega) for 15 mins at 37°C, phenol-chloroform extracted and ethanol precipitated, and then separated on a 5% polyacrylamide TBE-6M urea gel. Products were excised, crushed, and incubated in 500 μL RNA gel elution buffer (10 mM magnesium acetate, 0.5 M ammonium acetate, 1 mM EDTA) for 16 hours at 4°C. RNA was extracted from the eluate using phenol-chloroform extraction and ethanol precipitation. Approximately 50 pM of vigR 3’ UTR RNA was dephosphorylated using quick calf intestinal alkaline phosphatase (CIP, Thermo), then extracted using phenol-chlorofom and ethanol precipitation. The 5’ ends were radiolabelled with 20 μCi γ^32^P-ATP using T4 polynucleotide kinase (NEB) and separated from free nucleotides using a MicroSpin G-50 column (Cytiva), and then purified on denaturing PAGE as above. To analyse *vigR* 3’ UTR binding to full-length or sub-fragments of *dapE,* increasing excess amounts of the *dapE* RNAs were annealed to 50 fM of radiolabelled *vigR* 3’ UTR in 1x duplex buffer (40 mM Tris-acetate, 0.5 mM magnesium acetate, 100 mM NaCl) in a 10 μL reaction. These were incubated at 95°C for 5 minutes, then at 37°C for 2 hours. Samples were run on a 4% polyacrylamide 0.5X TBE gel containing 5% glycerol for ∼4 h at a maximum of 16V/cm or 1.33 mA/cm. Gels were then dried and visualised using a Fuji BAS-MP 2040 phosphorscreen and Typhoon FLA9500. Where appropriate 1.25 μM of antisense competitor oligonucleotides (**Supplementary Table 5**) were added to compete away radiolabeled *vigR* 3’ UTR at a concentration excess of 500x. RNA was annealed, run and visualised as above.

### Quantitative real-time PCR (qRT-PCR)

JKD6009 pICS3 or pICS3::*vigR* (Mediati et al., 2022) overnight cultures were diluted 1:100 into 10 mL fresh liquid BHI and grown at 37°C with 200 rpm shaking to OD_600nm_ 3.0. Cells were harvested by spinning at 3,800 *g* for 10 min at 4°C. A total of 5 U of recombinant RNasin (Promega) and 10 U of RQ1 RNase-free DNase (Promega) was added and RNA purified using the GTC-phenol:chloroform extraction procedure as previously described for *S. aureus* in Mediati et al., 2022. At least 1 μg of RNA was reverse-transcribed using SuperScript IV (Thermo). qRT-PCR was performed on a RotorGene Q (Eppendorf) using SensiFAST SYBR Hi-ROX (Bioline). A total cDNA concentration of 100 ng in combination with 200 nM of *dapE* oligonucleotide per reaction (**Supplementary Table 5**) resulted in ideal Ct values of between 8-12. The Ct values per reaction were calculated using the RotorGene Q analysis software (Qiagen). Relative gene expression was determined using ΔΔCt abundance of the *gapA* (SAA6008_RS08745, glyceraldehyde-3-phosphate dehydrogenase) transcript as a reference control.

### Galleria mellonella infection assay

The *G. mellonella* infection assay was performed as previously described (Frei et al., 2021). Briefly, *G. mellonella* larvae (230-250 mg) were injected with 10^7^ bacterial cells of each *S. aureus* construct (JKD6009, JKD6008 (isogenic parent), *vigR*^Δ3’UTR^, *vigR*^Δ3’UTR^-repair and Δ*isaA*) and a PBS control into the last right proleg using a 100 µL syringe (Hamilton Ltd). PBS-injected larvae resulted in no killing. The assay was done with 4 replicates using 5 larvae per replicate (*n*=20). Following infection, the larvae were incubated at 37°C for 6 consecutive days and monitored every 24 h for health and survival according to the *G. mellonella* Health Index Scoring System (Tsai et al., 2016). To examine vancomycin tolerance in JKD6009, JKD6008 and *vigR*^Δ3’UTR^ strains, larvae (230-250 mg) were infected with 10^7^ bacterial cells and incubated at 37°C for 1 h. Either vancomycin (10 mg/kg) or distilled water was injected into the infected larvae and treated as above. The assay was performed with 2 replicates using 10 larvae per treatment for each replicate (*n*=20).

## ACKNOWLEDGEMENTS

The pICS3 vector was a generous gift from Brice Felden (Université de Rennes). The authors thank Ian Monk for insightful discussions of *S. aureus* phylogenetics. DGM and JJT are supported by grants from the National Health and Medical Research Council (NHMRC, GNT1139313) and the Australian Research Council (ARC, DP220101938). This project received funding from the European Union’s H2020 research and innovation programme under Grant Agreement No. 753137. This work of the Interdisciplinary Thematic Institute IMCBio, as part of the ITI 2021-2028 program of the University of Strasbourg, CNRS and Inserm, was supported by IdEx Unistra (ANR-10-IDEX-0002), and by SFRI-STRAT’US project (ANR 20-SFRI-0012) and EUR IMCBio (ANR-17-EURE-0023) under the framework of the French Investments for the Future Program.

