## Supplementary Figures for "The 3’ UTR of *vigR* is required for virulence in *Staphylococcus aureus* and has expanded through STAR sequence repeat insertions"

### Supplementary Figure 1

**A**

```
               1       10        20        30        40        50        60        70        80        90       99
               |--------+---------+---------+---------+---------+---------+---------+---------+---------+--------|
     Mu50
  JKD6004      GATGTGAGTAGGACAGAAATGATAAGGGAGTG
  JKD6008      GAAGATTTAAAAAAGAATAAAAATGCATTATCTGATAAAGAATAAGATGTGAGTAGGACAGAAATGATAAGGGAGTGGGACAGAAATGATAAGGGAGTG
   USA300      GATGTGAGTAGGACAGAAATGATAAGGGAGTG
      COL      GATGTGAGTAGGACAGAAATGATAAGGGAGTG
Consensus      ....................................................................a.g.gagt.ggacagaaatgataagggagtg

               100     109       119       129       139       149       159       169       179       189      198
               |--------+---------+---------+---------+---------+---------+---------+---------+---------+--------|
     Mu50      GAT
  JKD6004      GGACAGAAATCATATTTTAGCAAAATTTATTTCGTCGTCCCACCCCGGCAAGGTTGACTAGAATTGAAAAAAGCTTGTTACAAGCGTATTTTCATTCAG
  JKD6008      GGACAGAAATCATATTTTAGCAAAATTTATTTCGTCGTCCCACCCCGGCAAGGTTGACTAGAATTGAAAAAAGCTTGTTACAAGCGTATTTTCATTCAG
   USA300      GGACAGAAATCATATTTTAGCAAAATTTATTTCGTCGTCCCACCCCGGCAAGGTTGACTAGAATTGAAAAAAGCTTGTTACAAGCGTATTTTCATTCAG
      COL      GGACAGAAATCATATTTTAGCAAAATTTATTTCGTCGTCCCACCCCGGCAAGGTTGACTAGAATTGAAAAAAGCTTGTTACAAGCGTATTTTCATTCAG
Consensus      ggacagaaatcatattttagcaaaatttatttcgtcgtcccaccccggcaaggttgactagaattgaaaaaagcttgttacaagcgtattttcattcAg

               199     208       218       228       238       248       258       268       278       288      297
               |--------+---------+---------+---------+---------+---------+---------+---------+---------+--------|
     Mu50      GTGAGTAGGACAGAAATGATAAAG-AACCACTATGATTTATTATATAGT----GGTTCTTTATCGTTAGTCATAGCTAATATGCATATAGAAAGAAGAA
  JKD6004      TCAACTACTGCCAATATAATATTGTAAACTATAGGACATTTATTAGTGTTTCAGGTTCTTTATCGTTAGTCATAGCTAATATGCATATAGAAAGAAGAA
  JKD6008      TCAACTACTGCCAATATAATATTGTAAACTATAGGACATTTATTAGTGTTTCAGGTTCTTTATCGTTAGTCATAGCTAATATGCATATAGAAAGAAGAA
   USA300      TCAACTACTGCCAATATAATATTGTAAACTATAGGACATTTATTAGTGTTTCAGGTTCTTTATCGTTAGTCATAGCTAATATGCATATAGAAAGAAGAA
      COL      TCAACTACTGCCAATATAATATTGTAAACTATAGGACATTTATTAGTGTTTCAGGTTCTTTATCGTTAGTCATAGCTAATATGCATATAGAAAGAAGAA
Consensus      tcaAcTActgCcaAtATaATAttGtAAaCtaTAgGAcaTtTatTAgtGTttcaGGTTCTTTATCGTTAGTCATAGCTAATATGCATATAGAAAGAAGAA

               298     307       317       327       337       347       357       367       377       387      396
               |--------+---------+---------+---------+---------+---------+---------+---------+---------+--------|
     Mu50      TACATGCGTGAAACTCATTCATAAGAAATACTAATATCTAAAGAAAACGTATTTCTTTATCTTTTCGAAATTCTCTGTGTTGGGGCCCTTACCCCAACC
  JKD6004      TACATGCGTGAAACTCATTCATAAGAAATACTAATATCTAAAGAAAACGTATTTCTTTATCTTTTCGAAATTCTCTGTGTTGGGGCCCTTACCCCAACT
  JKD6008      TACATGCGTGAAACTCATTCATAAGAAATACTAATATCTAAAGAAAACGTATTTCTTTATCTTTTCGAAATTCTCTGTGTTGGGGCCCTTACCCCAACT
   USA300      TACATGCGTGAAACTCATTCATAAGAAATACTAATATCTAAAGAAAACGTATTTCTTTATCTTTTCGAAATTCTCTGTGTTGGGGCCCTTACCCCAACT
      COL      TACATGCGTGAAACTCATTCATAAGAAATACTAATATCTAAAGAAAACGTATTTCTTTATCTTTTCGAAATTCTCTGTGTTGGGGCCCTTACCCCAACT
Consensus      TACATGCGTGAAACTCATTCATAAGAAATACTAATATCTAAAGAAAACGTATTTCTTTATCTTTTCGAAATTCTCTGTGTTGGGGCCCTTACCCCAACt

               397     406       416       426       436       446       456       466       476       486      495
               |--------+---------+---------+---------+---------+---------+---------+---------+---------+--------|
     Mu50      TGCACATTATTGAAAGCTG--------------------------------------------------------------------------------
  JKD6004      TGCACATTATTGAAAGCT---------------------------------------------------------------------------------
  JKD6008      TGCACATTATTGAAAGCT---------------------------------------------------------------------------------
   USA300      TGCACATTATTGAAAGCTGACTATTGGCCAGCTTCTGTGTTGGGGCCC---------------------------------------------------
      COL      TGCACATTATTGAAAGCTGACTATTGGCCAGCTTCTGTGTTGGGGCCCCGCCAACTTGCACATTATTGAAAGCTGACTATTGGCCAGCTTCTGTGTTGG
Consensus      TGCACATTATTGAAAGCTg................................................................................

               496     505       515       525       535       545       555       565       575       585      594
               |--------+---------+---------+---------+---------+---------+---------+---------+---------+--------|
     Mu50      ---------------------------------------------------------------------------------------------------
  JKD6004      ---------------------------------------------------------------------------------------------------
  JKD6008      ---------------------------------------------------------------------------------------------------
   USA300      ---------------------------------------------------------------------------------------------------
      COL      GGCCCCGCCAACTTGCATTGTCTGTAGAATTTCTTTTCGAAATTCTCTGTGTTGGGGCCCTTACCCCAACTTGCACATTATTGAAAGCTGACTATTGGC
Consensus      ...................................................................................................

               595     604       614       624       634       644       654       664       674       684      693
               |--------+---------+---------+---------+---------+---------+---------+---------+---------+--------|
     Mu50      --------------------------------------------------ACTATTGGCCAGCTTCTGTGTTGGGGCCCCGCCAACTTGCATTGTCTGT
  JKD6004      -------------------------------------------------AACTATTGGCCAGCTTCTATGTTGGGGCCCCGCCAACTTGCATTGTCTGT
  JKD6008      -------------------------------------------------AACTATTGGCCAGCTTCTATGTTGGGGCCCCGCCAACTTGCATTGTCTGT
   USA300      --------------------TTACCCCAACTTGCACATTATTGAAAGCTAACTATTGGCCAGCTTCTATGTTGGGGCCCCGCCAACTTGCATTGTCTGT
      COL      CAGCTTCTGTGTTGGGGCCCTTACCCCAACTTGCACATTATTGAAAGCTAACTATTGGCCAGCTTCTATGTTGGGGCCCCGCCAACTTGCATTGTCTGT
Consensus      .................................................aACTATTGGCCAGCTTCTaTGTTGGGGCCCCGCCAACTTGCATTGTCTGT

               694     703       713       723       733       743       753       763       773       783      792
               |--------+---------+---------+---------+---------+---------+---------+---------+---------+--------|
     Mu50      AGAATTTCTTTTCGAAATTCTCTGTGTTGGGGCCCCGGGGCGCATTTTCGTTCGGTTAATTACTGTCAATATTAAATTGTGTGCCTAGTACATTGACTT
  JKD6004      AGAATTTCTTTTCGAAATTCTCTATGTTGGGGCCCCGGGGCGCATTTTCGTTCGGTTAATTACTGTCAATATTAAATTGTGTGCCTAGTACATTGACTT
  JKD6008      AGAATTTCTTTTCGAAATTCTCTATGTTGGGGCCCCGGGGCGCATTTTCGTTCGGTTAATTACTGTCAATATTAAATTGTGTGCCTAGTACATTGACTT
   USA300      AGAATTTCTTTTCGAAATTCTCTATGTTGGGGCCCCGGGGCGCATTTTCGTTCGGTTAATTACTGTCAATATTAAATTGTGTGCCTAGTACATTGACTT
      COL      AGAATTTCTTTTCGAAATTCTCTATGTTGGGGCCCCGGGGCGCATTTTCGTTCGGTTAATTACTGTCAATATTAAATTGTGTGCCTAGTACATTGACTT
Consensus      AGAATTTCTTTTCGAAATTCTCTaTGTTGGGGCCCCGGGGCGCATTTTCGTTCGGTTAATTACTGTCAATATTAAATTGTGTGCCTAGTACATTGACTT

               793     802       812       822       832       842       852       862       872       882   888
               |--------+---------+---------+---------+---------+---------+---------+---------+---------+-----|
     Mu50      ATATCTCGAGCTCTTGAATATGTGTATTTTAAATATATAAATTAAAAAAGGAACAAAGCAGCTATTTATGATTTGTGTCATATCAATAGCATCGTT
  JKD6004      ATATCTCGAGCTCTTGAATATGTGTATTTTAAATATATAAATTAAAAAAGGAACAAAGCAGCTATTTATGATTTGTGTCATATCAATAGCATCGTT
  JKD6008      ATATCTCGAGCTCTTGAATATGTGTATTTTAAATATATAAATTAAAAAAGGAACAAAGCAGCTATTTATGATTTGTGTCATATCAATAGCATCGTT
   USA300      ATATCTCGAGCTCTTGAATATGTGTATTTTAAATATATAAATTAAAAAAGGAACAAAGCAGCTATTTATGATTTGTGTCATATCAATAGCATCGTT
      COL      ATATCTCGAGCTCTTGAATATGTGTATTTTAAATATATAAATTAAAAAAGGAACAAAGCAGCTATTTATGATTTGTGTCATATCAATAGCATCGTT
Consensus      ATATCTCGAGCTCTTGAATATGTGTATTTTAAATATATAAATTAAAAAAGGAACAAAGCAGCTATTTATGATTTGTGTCATATCAATAGCATCGTT
```

#### Supplementary Figure 2

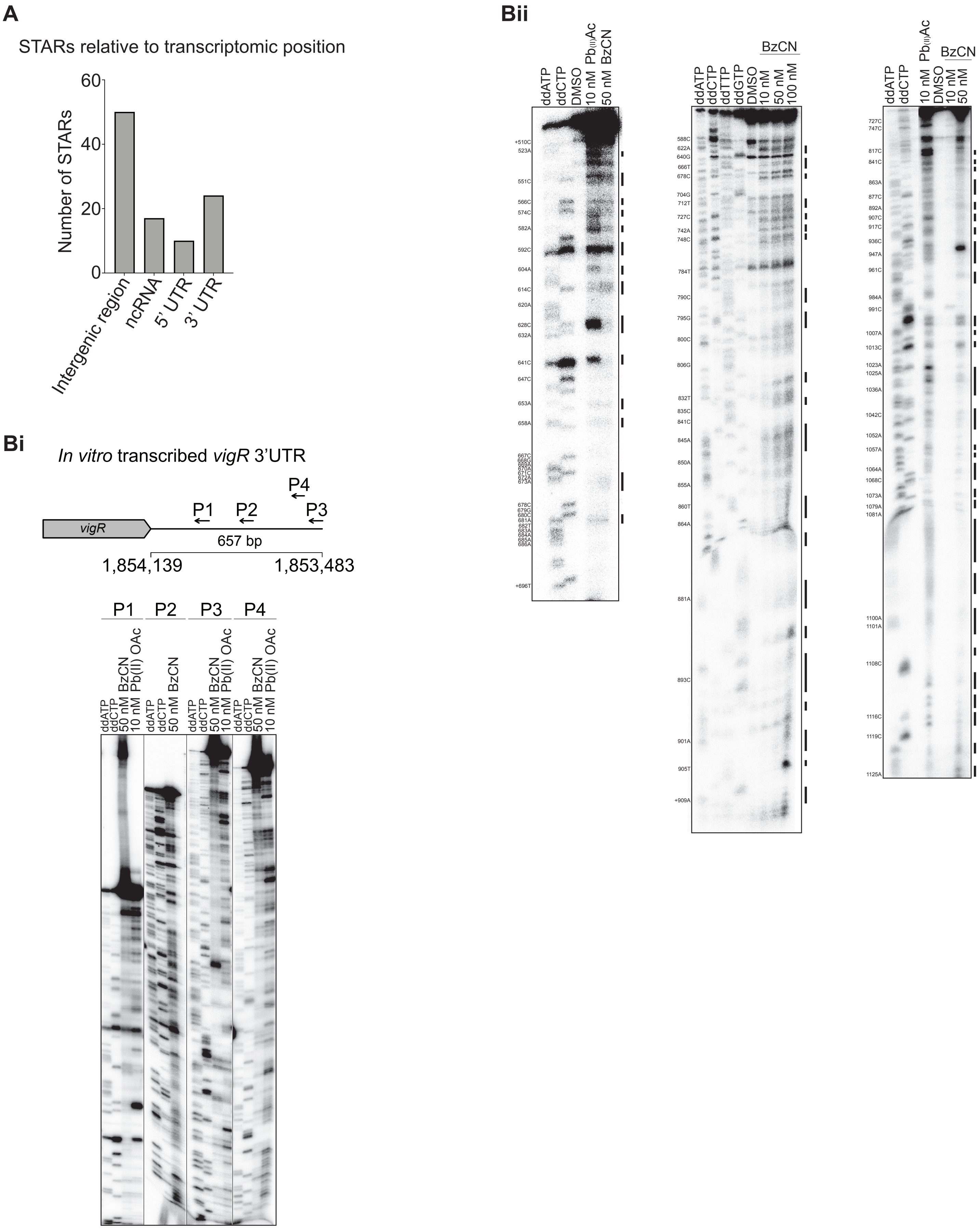

**Supplementary Figure 3**

**A**

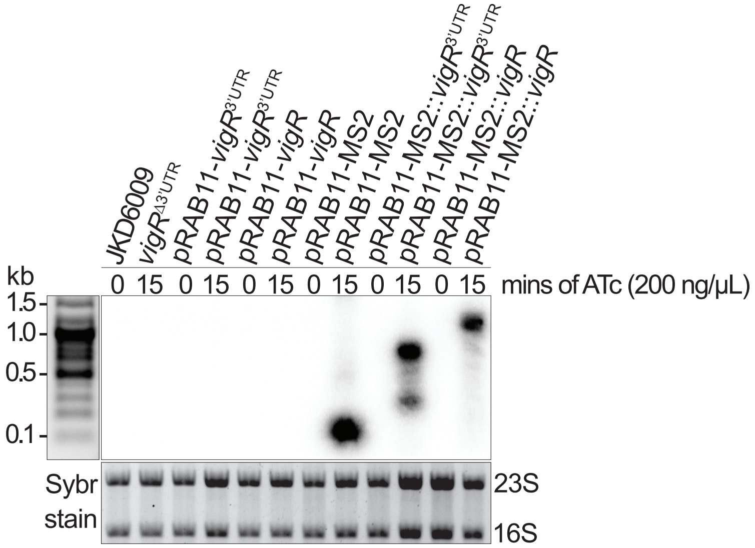


**B**

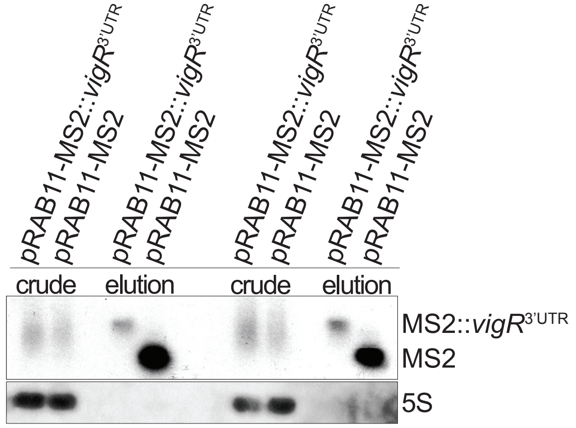

#### Supplementary Figure 4

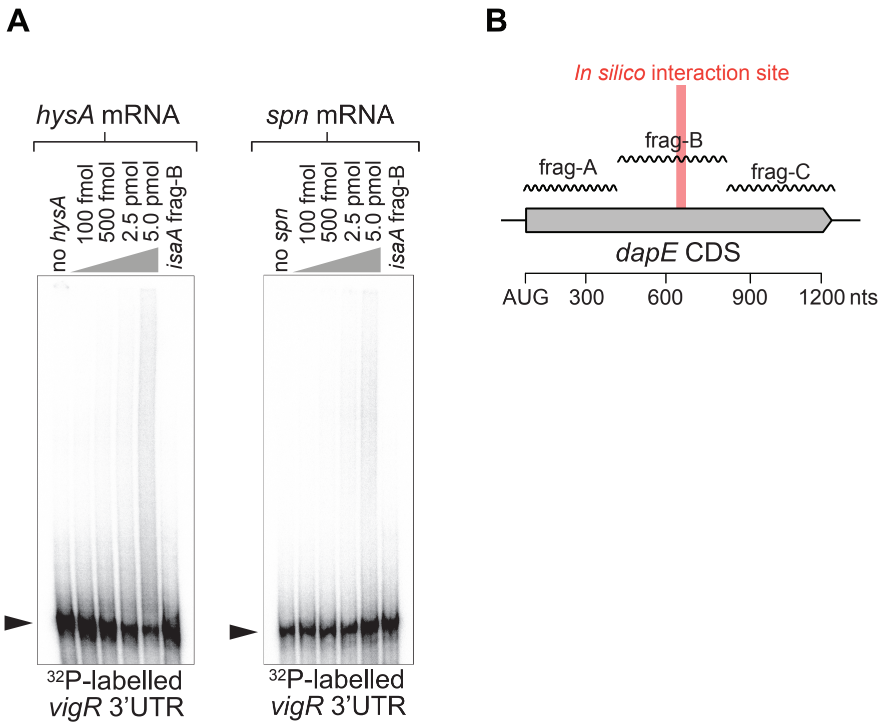

**Supplementary Figure 5**

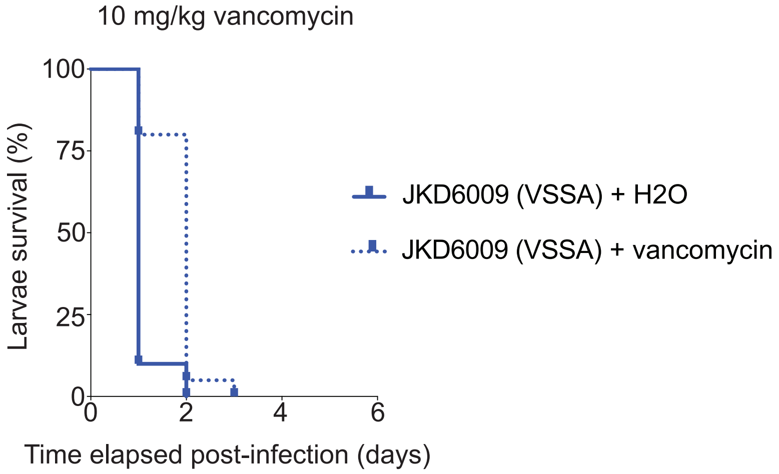
